# Genome-wide fitness profiling reveals limited modeled microgravity-specific gene requirements in *Vibrio fischeri*

**DOI:** 10.1101/2020.08.13.250431

**Authors:** Emanuel Burgos, Madeline M. Vroom, Ella Rotman, Megan Murphy-Belcaster, Jamie S. Foster, Mark J. Mandel

**Affiliations:** Department of Medical Microbiology and Immunology, University of Wisconsin-Madison, Madison, WI USA; Department of Microbiology and Cell Science, Space Life Science Lab, University of Florida, Merritt Island, FL USA; Department of Microbiology-Immunology, Northwestern University Feinberg School of Medicine, Chicago, IL USA

**Keywords:** modeled microgravity, essential genes, symbiosis, *Vibrio fischeri*, *Aliivibrio fischeri*, *Euprymna scolopes*, INSeq, Tn-seq

## Abstract

To support long-duration spaceflight, identifying genes that enable microbial survival and stability under space stressors is crucial, as these factors can affect both host health and in-space biomanufacturing efforts. Our objective was to determine what bacterial genes are required for growth in culture under modeled, or simulated, microgravity conditions compared to normal gravity controls. We focused on the marine bacterium *Vibrio fischeri*, which forms a monospecific symbiosis with the Hawaiian bobtail squid, *Euprymna scolopes*. This symbiotic microbe has been studied during spaceflight and in ground-based modeled microgravity conditions and is emerging as a valuable model organism for in-space biomanufacturing. Using transposon insertion sequencing (INSeq), we identified dozens of genes that exhibited fitness defects under both conditions, yet we identified relatively few genes with differential effects under modeled microgravity or gravity specifically. We additionally compared RNA-seq and INSeq data and determined that expression under simulated microgravity was not predictive of the essentiality of a given gene. In summary, empirical determination of conditional gene essentiality identifies few microgravity-specific genes for environmental growth of *V. fischeri*, suggesting that the condition of microgravity has a minimal impact on gene requirement during growth in media. These findings suggest that engineered strains of *V. fischeri* for on-demand biomanufacturing applications may require minimal genetic adaptation during spaceflight, easing the challenges of maintaining robust microbial production platforms for long-duration space missions.

## INTRODUCTION

Spaceflight alters microbial physiology and as the number and duration of crewed space missions expands beyond low Earth orbit, elucidating the mechanisms underlying these changes is essential^1–10^. Although previous studies using both natural and modeled microgravity conditions have resulted in a wide range of microbial physiological and genetic responses, many initial studies primarily targeted pathogenic strains^11–13^. More recently, studies have begun to examine the effects of microgravity on beneficial microbes that associate with animals and how spaceflight impacts these interactions^4–6,10,14,15^, as well as engineered microbes that serve as platforms for the in-space biomanufacturing of essential bioproducts during space exploration^16,17^. Given the importance of microbes on host health and the biomanufacturing of high-value products during spaceflight and, we sought here to ask a fundamental question of what genes are required for bacterial growth in microgravity.

Due to the number of logistical constraints in conducting spaceflight experiments, several ground-based platforms have been developed to model key features of the microgravity environment^18^. Rotating cell culture systems equipped with High Aspect-Ratio Vessels (HARVs) are widely used as they generate a low-shear, low-turbulence fluid environment that approximates aspects of microbial growth experienced in low Earth orbit and have been used for decades to study biological responses to modeled microgravity^19^. As cells grow in the HARVs, the hydrodynamic forces within each vessel offset the effects of gravity and the cells are essentially in “freefall” and maintained in a constant suspension (**Fig. 1**). This environment, commonly referred to as low-shear modeled microgravity (LSMMG), can induce diverse microbial physiological responses, including altered growth, virulence, biofilm formation, secondary metabolite production, stress responses, and antibiotic susceptibility^6,20–25^. However, the magnitude of these responses can vary considerably among microbial taxa and strains and are influenced by experimental parameters, such as growth medium, temperature, nutrient availability, culture duration, and environmental stressors^20–25^. Despite these potential caveats and limitations using platforms, such as HARVs, to simulate the microgravity environment, identifying genes that are required for bacterial survival and growth under LSMMG conditions represents an important step toward defining the core genetic determinants that enable microorganisms to adapt to spaceflight-like environments.

**Figure 1.**
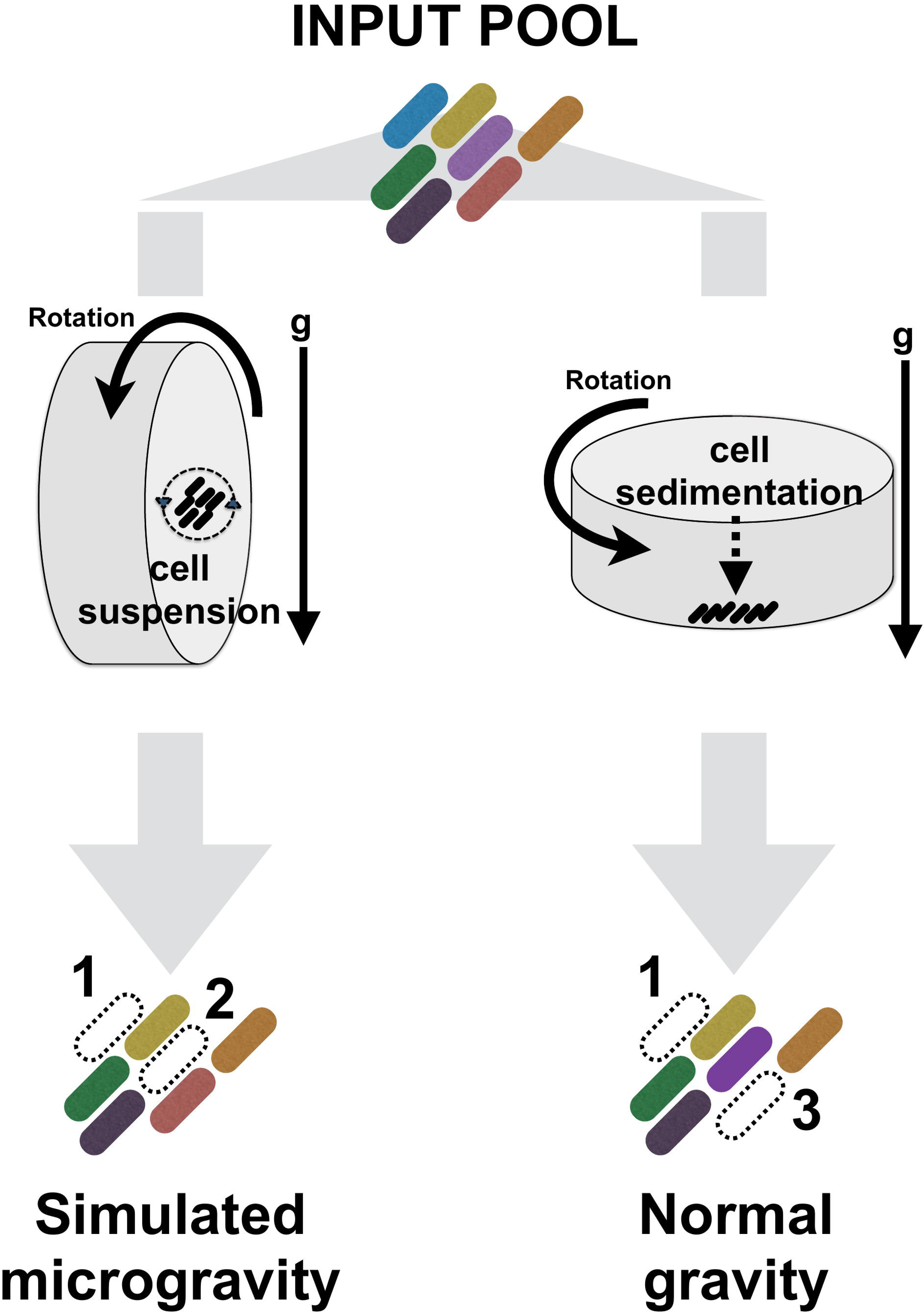
Overview of the impact of low-shear modeled microgravity (LSMMG) on cells in High Aspect Ratio Vessels (HARVs) compared to gravity controls. Cartoon depicting the experimental setup for bacterial mutant enrichment in HARV vessels to model microgravity (LSMMG) and gravity controls. INSeq libraries of *Vibrio fischeri* were grown in HARV vessels. Mutants exhibited either no defects in either condition or exhibited (1) a growth defect in both conditions, (2) LSMMG defect, (3) or a gravity-specific defect.

One microbe that has emerged as a versatile model to explore the impact of space-like stressors on microbial physiology is the marine bacterium *Vibrio fischeri*, which forms a mutualistic association with the Hawaiian bobtail squid, *Euprymna scolopes* and has been studied extensively under both modeled and actual spaceflight conditions^1,10,26,27^. The bacteria colonize the dedicated symbiotic light organ of the host, where bioluminescence from the bacteria is projected downward and camouflages the host in the moonlight^28^. This system has been especially valuable to understand the molecular basis by which animals acquire specific microbes from the environment^29^. The bacteria are genetically manipulatable, and the site of colonization can be imaged directly in live animals, enabling studies that have revealed much of the molecular dialogue between the two partners^30–33^. The specificity of the relationship, in which only certain strains of *V. fischeri* can colonize the squid host, provides a strong model system to study animal-microbiome formation, development, and evolution^34^. Furthermore, the small size of the host animal and simplicity of the symbiosis has contributed to the value of this system for studying host-microbe interactions during spaceflight^1,3,10,35^.

In this study, we take a complementary approach to previous work on bacterial gene expression under simulated microgravity. Previous studies using the squid-vibrio system have focused extensively on the impact of simulated and actual spaceflight conditions on the host squid physiology in the presence and absence of *V. fischeri*^1,3,10,35,36^. In these previous studies, results have shown that under both simulated and actual spaceflight, the colonization of the host by *V. fischeri* results in the down regulation of several host genes associated with oxidative and heat stress, suggesting that being in a symbiotic state is beneficial to the host during spaceflight^3,10^. These studies do not, however, examine the impact of the space-like environment on the symbiont, specifically, what genes are required for survival under microgravity and modeled microgravity conditions. Other studies in the symbiont have applied a genome-scale genetic approach technology to globally identify *V. fischeri* genes that are essential, and conditionally essential, under specific environmental conditions and during growth in media under laboratory conditions^37^. Here, we apply a similar global approach to determine what genes are required for bacterial growth under simulated microgravity conditions to generate a baseline for *V. fischeri* cultures on the impact that microgravity-like conditions have on symbiont health and physiology.

## RESULTS

### Global analysis of bacterial genes required for growth in low-shear modeled microgravity

Previous studies have shown that under LSMMG conditions there is a significant increase in growth rates of bacteria compared to gravity controls, including the marine bacterium *V. fischeri*^6,38,39^, suggesting that the LSMMG environment within the HARVs provides a physiological advantage to the cells. Additionally, a number of studies have examined differential gene expression in beneficial microbes under microgravity or simulated microgravity conditions^40–43^, including *V. fischeri*^6^. The current study asks a distinct, yet related, question of what genes are differentially required for growth under simulated microgravity conditions compared to a gravity control condition.

To identify mutants with a competitive growth defect in simulated microgravity, we applied the transposon insertion sequencing (INSeq/Tn-seq) approach to *V. fischeri* strains cultivated under HARV conditions in both the low-shear modeled microgravity (LSMMG) and gravity positions (**Fig. 1**)^44,45^. We started with a characterized mutant library of over 40,000 transposon mutants in *V. fischeri* strain ES114^37^. The resulting “input” library was introduced into HARV vessels under either LSMMG or normal gravity conditions (1 x *g*). No significant growth difference in the HARV was observed compared to previous studies (Fig. S1) ^6,38,39^. We simultaneously examined the mutant library grown in LSMMG and gravity control conditions after growth for approximately 15 generations, and each resulting “output” pool of mutants was frozen. A total of six LSMMG biological replicates and three gravity control replicates were compared to the input library. From each sample, DNA was isolated, from which INSeq libraries were constructed^45^. The libraries were sequenced on an Illumina HiSeq 2000 instrument, and the location of each transposon insertion was mapped using the pyinseq Python package.

For each sample, we obtained between 6.65 - 8.18 x 10^6^ Illumina reads (**Table S1**). We observed similar numbers and patterns of unique transposon hits across the samples tested, suggesting that the library did not undergo significant bottlenecks during the experiment (**Table S1**). Next, for each sample, the normalized transposon insertion counts (CPM; counts-per-million Illumina reads) was examined in each gene. We compared the similarity of these gene-level counts across the samples in the analysis using pairwise correlation analysis. All pairwise comparisons had a high level of similarity (Spearman R^2^ > 0.95), and the HARV-grown samples were clearly distinguishable from the input libraries (**Fig. 2**). Examination of the heat map suggested little overall differentiation between the LSMMG and gravity samples that were otherwise grown similarly. We proceeded to compare individual genes that were depleted under LSMMG, gravity, or both conditions.

**Fig. 2.**
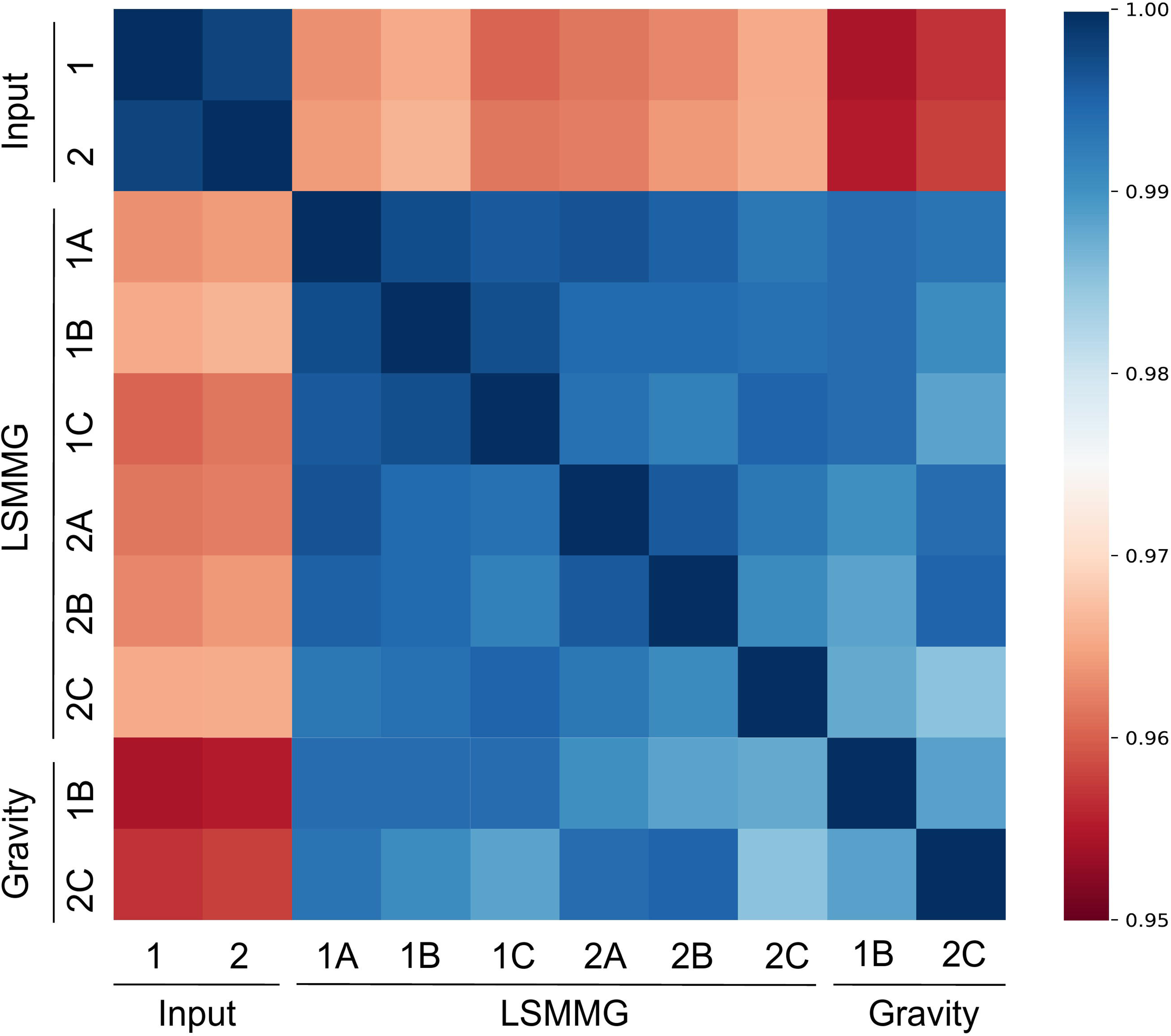
Pairwise correlation between samples revealed a clear difference between input samples and output samples from the HARV conditions. Spearman correlations between samples, in which the normalized transposon insertion counts in each gene were compared. Input replicates were the original INSeq library grown for inoculation into the HARV, and low-shear modeled microgravity (LSMMG) and normal gravity replicates were outputs from the respective HARV-grown experiments. All pairwise comparisons had a minimum Spearman ^2^ > 0.95.

To identify genes that had significant differential depletion under modeled microgravity we used DESeq2 to calculate the median representation of each mutant in the output LSMMG or gravity pools, and plotted those values compared to the input pool. We focused our analysis on genes that were depleted at least 2-fold in the different conditions. Genes that did not meet these criteria, genes that were poorly represented in the input pool, and genes for which a previous study suggested that mutants impaired bacterial growth were excluded from this analysis (**Fig. 3**; open black circles). A total of 109 genes exhibited depletion under both LSMMG and gravity conditions (**Fig. 3**; filled circles). Of these genes, most were similarly depleted under both conditions. However, there were genes in this group that had a ≥ 2-fold difference in one condition (LSMMG or gravity) relative to the other, including 10 genes more depleted under gravity conditions and one gene (*rodA*) more depleted under LSMMG. Furthermore, there were two genes in the analysis that were significantly depleted (p < 0.05 from DESeq2 analysis) under gravity conditions and not under LSMMG conditions (*flgD, rfaD*). There were no genes that were only significantly depleted under LSMMG conditions.

**Fig. 3.**
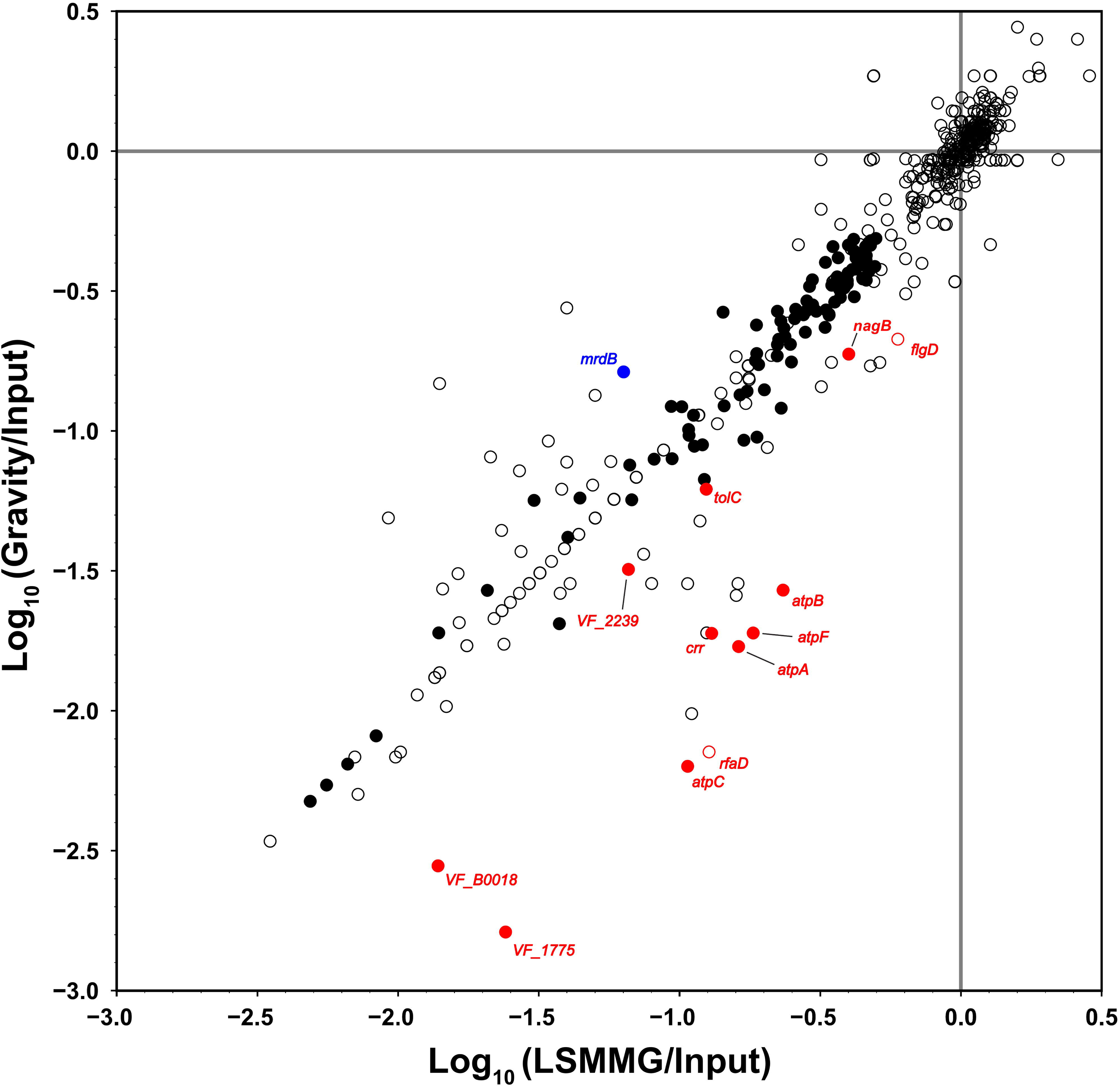
Mutant behavior under LSMMG or gravity revealed fitness in each HARV condition, normalized to the input transposon sequencing library. A) Comparison scatter plot of microgravity (x-axis) and gravity (y-axis) Log_10_(Fold-Change) values. Counts for each gene for each replicate were normalized and used to calculate Log_10_(Fold-Change) values as described in Materials and Methods. Genes that were poorly represented in the input pool, did not exhibit significant depletion, or were previously determined to be growth deficient in LBS medium were designated as open black circles. Filled black circles are genes that were significantly (corrected False Discovery Rate p-value < 0.05) depleted 2-fold change under both simulated microgravity and gravity conditions. Genes that are filled red were significant in both conditions but more depleted under gravity, whereas open red circles are exclusively significant in gravity. Filled blue genes are significant in both, but more depleted under simulated microgravity.

Given the differential sample sizes and the small effects, we next compared the two output conditions directly, taking advantage of the robust statistical tests within DESeq2 to identify genes that were significant between the two output conditions even if they were not significant between an individual output condition and the input pool. The volcano plots shown in **Fig. 4** show the results of this analysis, with raw data in **Table S2**. While each of the HARV output conditions (gravity or LSMMG) exhibited >400 genes that were significantly depleted relative to the input pool, direct comparison of the two output conditions revealed only 13 genes with statistically significant differences, and only two with a fold change of 2.0-2.4 (*nagB, spoT*; **Fig. 4C, Table S2**), compared to effects exceeding a 250-fold change in the comparisons to the input sample (**Fig. 4A, 4B**). Therefore, the INSeq data suggest that the effect of the HARV environment broadly had a much greater impact on gene requirement than the gravity condition within the HARV.

**Fig. 4.**
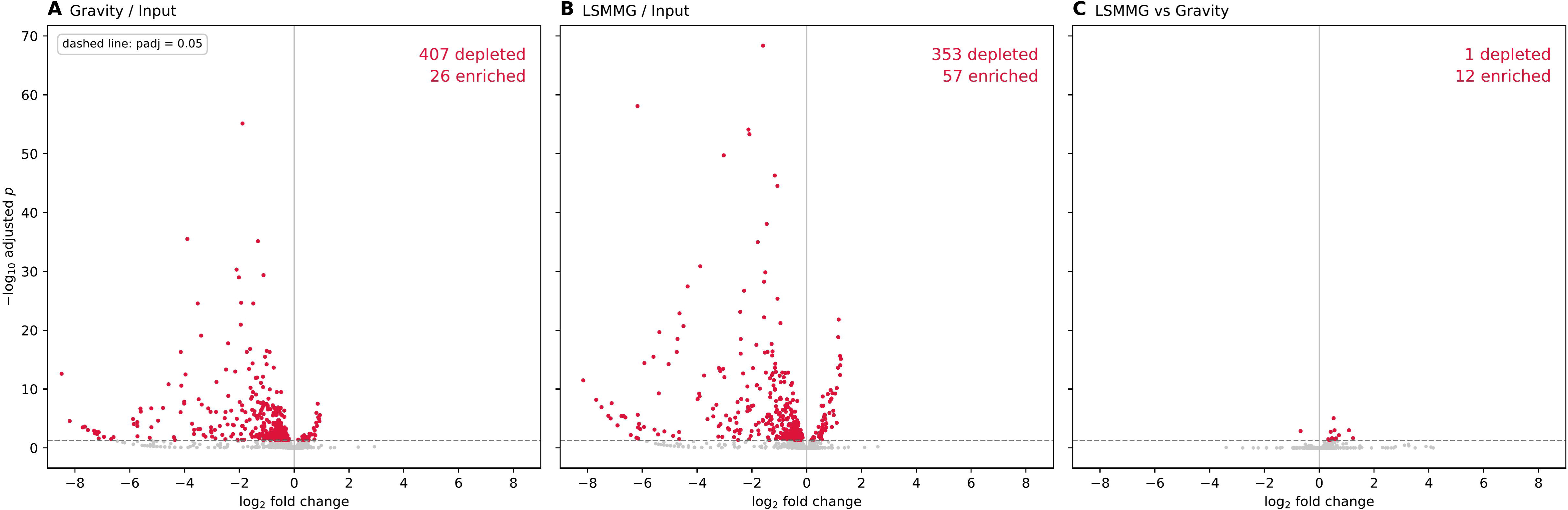
Volcano plots showing direct comparison of INSeq data across conditions. (A) Mutants depleted in Gravity vs Input, (B) Mutants depleted in LSMMG vs Input, (C) Mutants depleted in LSMMG vs Gravity, as assessed by DESeq2. Red dots are statistically significant (*p_adj_* < 0.05). Raw data are in Table S2.

### Validation of INSeq results through defined mutant competitions

The above results were derived from an analysis of complex mutant pools with > 40,000 mutants. We therefore sought to determine whether we would observe the same behavior using one-versus-one competitions between defined mutant strains and the parental strain. For this analysis, we considered the entire data set (i.e., **Table S3**), and we proceeded to isolate mutant strains that had a depletion from the INSeq analysis, as well as mutants in two control genes that were not depleted under either condition (*brnQ* and *nhaR*). We examined mutants that had a range of fitness defects and that were represented in a mutant collection in the laboratory. Each defined mutant strain was grown in culture, then competed in the HARVs against the parental strain that carries the LacZ-expressing plasmid pVSV103^46^. The input and output pools from each experiment were plated onto LBS-Xgal medium, and the ratios of blue:white colonies in the samples were calculated. A competitive index was calculated to determine the fitness of each mutant under each condition. The results of this analysis are plotted in **Fig. 5**. There was strong overall concordance between the INSeq and defined competition results. All the genes for which mutants were significantly depleted under INSeq were also substantially depleted in the defined competitions, and the control mutants exhibited no substantial depletion. Therefore, we conclude that the INSeq analysis can reliably predict conditional gene requirements in the simulated microgravity environment.

**Fig. 5.**
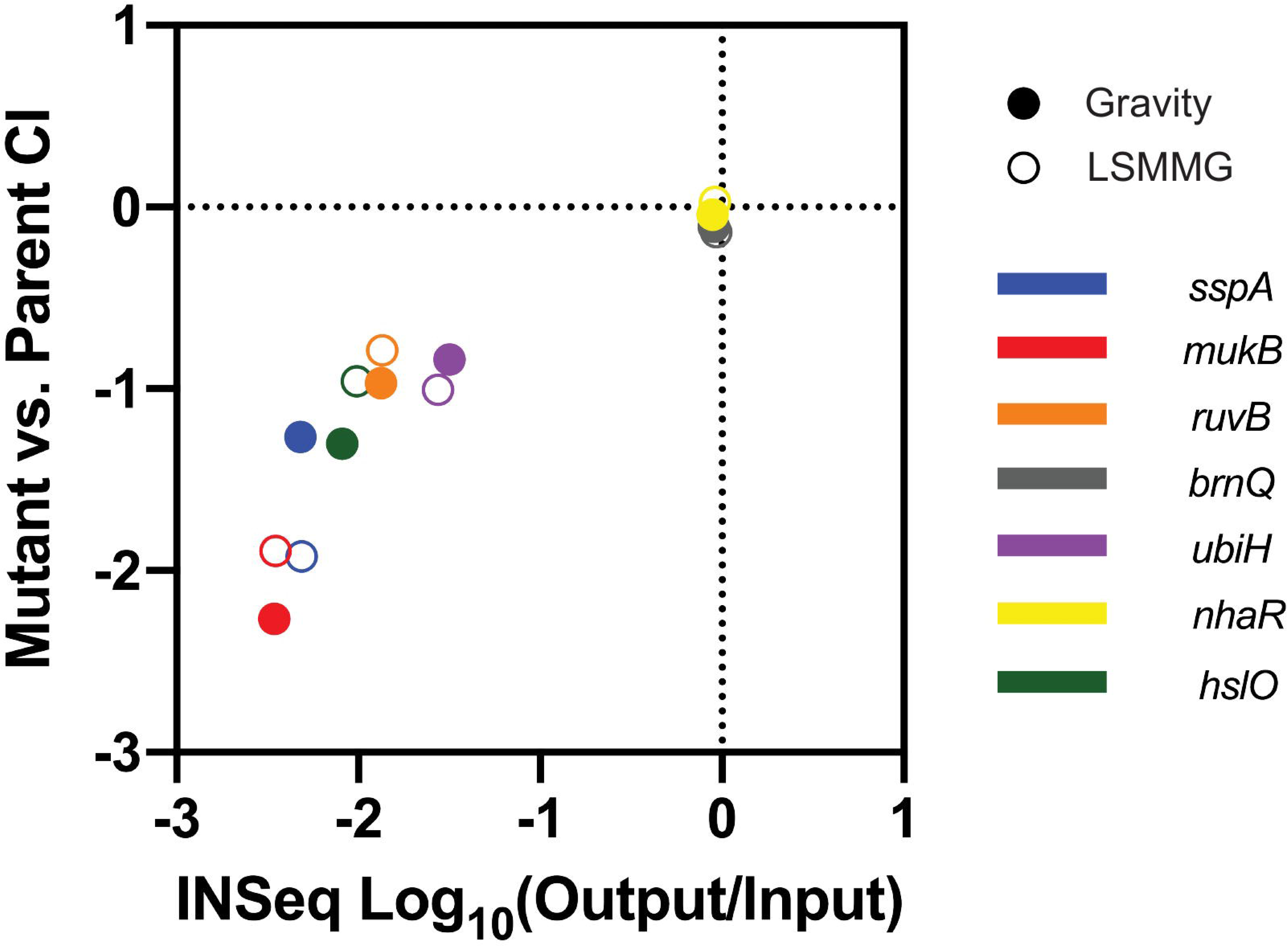
Competition of defined mutants provided validation of mutant fitness from within the complex INSeq library. Mutants were competed against wild-type *V. fischeri* under simulated microgravity and gravity using the HARV vessels. Dots represent fold-change for 1:1 competition of mutant versus marked parental strain (MJM1575).

### Examination of the relationship between bacterial gene requirement and gene induction in modeled microgravity

We recently examined bacterial genes for which mRNA levels are induced in the modeled microgravity condition, compared to gravity controls^6^. We analyzed our current study of gene requirement (i.e., INSeq) with the previous dataset on gene expression (i.e., RNA-Seq) to ask whether there is a correlation between gene requirement and gene expression. As shown in **Fig. 6A**, there was no significant correlation between gene requirement and transcript induction (Pearson *r* = 0.08) when compared to published transcriptome data from the HARV platform^6^. Our data above suggest that the HARV platform may play a more significant role in shaping mutant communities than the specific LSMMG or gravity conditions as has been determined for other bacterial strains^47^. Therefore, we explored whether genes required in the HARV are also induced in the HARV when normalized to non-HARV samples. For our INSeq data, we normalized to the input sample that did not experience the HARV condition, and for RNA-Seq, we normalized to published data on culture-grown *V. fischeri* ES114 on LBS medium^48^. As shown in **Fig. 6B-C**, there was a lack of overall correlation between the gene expression and gene requirement (Pearson *r* = -0.01, -0.08 for **Fig. 6B, 6C**, respectively). We validated the quality of the published RNA-seq datasets by confirming consistent expression of housekeeping transcripts across the datasets (e.g., *gyrB, recA, rpoA*). Therefore, we conclude that gene requirement in the HARV cannot be predicted from transcriptome induction data, emphasizing the need for empirical determination of gene essentiality.

**Fig. 6.**
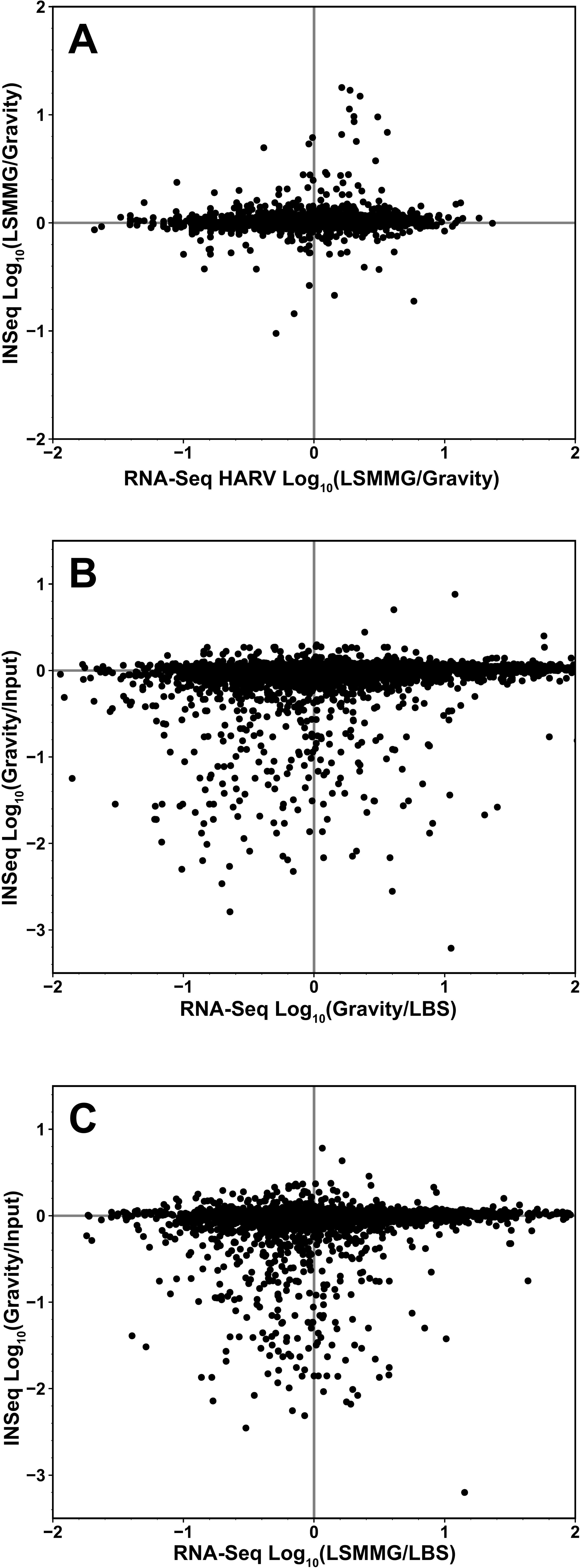
Gene expression (RNA-Seq) does not predict conditional gene requirement (INSeq) under LSMMG conditions. INSeq (y-axis) and RNA-Seq (x-axis) were compared by using log-normalized fold-change values for different experimental conditions. A) INSeq (LSMMG/Gravity) was compared to a previously published RNA-Seq (LSMMG/Gravity) under HARV conditions. We also calculated fold-changes for RNA-Seq HARV conditions using a dataset from Luria-Bertani media (LBS medium): B) INSeq (Gravity/Input) were compared to RNA-Seq (Gravity/LBS) and C) INSeq (LSMMG/Input) was compared to RNA-Seq (LSMMG/LBS). As shown in **Fig. 5A**, there was no significant correlation between gene requirement and transcript induction (Pearson *r* = 0.08). Our data above suggest that the HARV platform may play a more significant role in shaping mutant communities than the specific LSMMG or gravity conditions as has been determined for other bacterial strains^47^. Therefore, we explored whether genes required in the HARV are also induced in the HARV^6^ when normalized to non-HARV samples. For our INSeq data, we normalized to the input sample that did not experience the HARV condition, and for RNA-Seq, we normalized to published data on culture-grown *V. fischeri*^48^. As shown in **Fig. 5B-C**, there was a lack of overall correlation between the gene expression and gene requirement (Pearson *r* = -0.01, -0.08 for **Fig. 5B, 5C**, respectively). Therefore, we conclude that gene requirement in the HARV cannot be predicted from transcriptome induction data, emphasizing the need for empirical determination of gene essentiality.

## DISCUSSION

This study provides a global view of *V. fischeri* genes that are required for growth under simulated microgravity conditions. Given the prominence of modeled microgravity platforms, such as HARVs, in examining the impact of spaceflight on animal-microbe interactions, a motivation for this work was to understand how the basic requirements for bacterial survival and growth are altered during growth in the microgravity analog platforms. A surprising finding was that there was little difference in gene requirement under the LSMMG conditions compared to gravity controls, despite dramatic differences in gene expression. These findings, and their implications, are discussed in greater detail below.

The global data provided by INSeq, along with the comparison of the INSeq data with results from defined mutants, provided sensitive internal controls to examine growth in multiple HARV bioreactors. Overall, the data we obtained from HARVs were highly consistent regardless of the LSMMG or gravity orientation. As shown in the heat map in **Fig. 2**, the high correlation of INSeq replicates within a treatment (e.g., for LSMMG samples) suggests that there was no substantial bottleneck during inoculation or growth in the HARVs. Furthermore, a strong correlation between INSeq results and the results with defined mutants (**Fig. 5**) provides support that the global data are representative of gene-level data. In fact, the consistency we observed between vessel replicates was also observed between LSMMG and gravity samples in the HARVs. The heat map in **Fig. 2**, coupled with the tight correlation of the samples in **Fig. 3** (Pearson R^2^ = 0.96) and the direct output comparison in **Fig. 4**, illustrates that there was little variation observed between genes required for bacterial growth under both LSMMG and gravity conditions.

As *V. fischeri* is a genetically tractable bacterium capable of producing and exporting biologically active compounds for space biomanufacturing applications^38^, this result indicates there will not be a confounding effect of genes that are simply required for bacterial growth under microgravity. Put another way, in the future, if genes are identified that are required for optimizing target biomolecule production or host animal colonization under microgravity, they are likely to be required specifically for that interaction or function, and not simply for growth under this altered gravity condition.

A common follow-up experiment for INSeq experiments is to validate the global data with individual gene mutants. We conducted such an experiment in **Fig. 5** and found that the results were consistent across over two orders of magnitude. Another common follow-up experiment is to complement back the wild-type allele for mutants under study to then demonstrate that the wild-type phenotype is restored. Given the negative result that we observed—i.e., that mutants did not display different effects in simulated microgravity compared to the gravity control (e.g., **Fig. 4**)—there was not the impetus to conduct this experiment. However, as described above, we argue that this negative finding is an important one given the expectation of the experiment, and given the rigor with which we validated the results with the single-gene mutants in **Fig. 5**.

Despite the overall patterns of concordance between LSMMG and gravity, there were genes for which we observed differential effects between the two conditions, as plotted in **Fig. 3** and detailed in **Table S4**. For example, depletion in *rodA (mrdB)* under both LSMMG and gravity was observed, yet the gene was more depleted under LSMMG. RodA is a SEDS-family peptidoglycan polymerase that has multiple effects on bacterial cell shape and division ^49^. This result suggests that differences in bacterial morphology under the two different conditions may impact the genetic requirement for *rodA*, though we note that its absence does affect growth under both regimes.

We additionally identified several genes with mutants depleted under both conditions but that were more depleted under gravity (**Fig. 3**, filled red dots). Notably, multiple genes for the F0F1-ATPase were depleted under both conditions, but more so under gravity. Interestingly, prior work in *Escherichia coli* demonstrated that a similar set of genes was essential for aerobic growth in minimal glycerol media, even though the metabolic model used predicted that they would not be required^50^. Together with our results, this suggests that this subset of genes (*atpA, atpB, atpC, atpF*) may perform a function separate from their role in ATP synthesis. Finally, there were two genes for which mutants exhibited significant depletion under gravity but not LSMMG: *flgD* (encoding the hook capping protein FlgD) and *rfaD* (encoding the LPS biosynthesis enzyme ADP-L-glycero-D-mannoheptose 6-epimerase). Both genes affect the outer surface of the bacteria, and consistent with the *rodA* results above, support the idea that the bacterial envelope is most susceptible to differential effects of gravity.

It is also important to note that we observed dozens of genes for which mutants exhibited similar depletion under both LSMMG and gravity conditions (**Fig. 3**; black filled dots). We note that significant growth defects for mutants in these genes were not observed during 15 generations of growth in LBS medium under aerobic conditions^37^. Therefore, these genes are likely required for robust growth in the HARV, but not for growth in LSMMG versus gravity. Although the HARV environment is low shear, it is not oxygen limited as there is continuous oxygen diffusion through the gas permeable silicone membrane with estimated oxygen availability at 5750 µmol O_2_ per hour^51^. Despite the availability of O_2_, it is possible that the *V. fischeri* cells are experiencing changes in cellular respiratory demand, membrane energetics or issues maintaining ion gradients withing the HARVs^52^. This hypothesis is supported by the presence of four genes of Na^+^-translocating NADH:quinone oxidoreductase (*nqrA*, *nqrB*, *nqrD*, *nqrE*), deletion of which in the related species *V. cholerae* decreases respiratory Na⁺ extrusion and membrane-potential generation and causes broad metabolic changes^53–55^. We also note the presence of some genes that are required for robust symbiosis in this category, including *degS*, *dnaJ*, and *ompU*^37,56^. Given that mutants in these genes exhibited growth defects in both LSMMG and gravity conditions in the HARVs, our results suggest that colonization in the HARV may proceed differently than under standard laboratory conditions.

A possible limitation of the current study is that the transposon library used was built on agar plates in a standard microbiology laboratory (i.e., under normal gravity conditions). However, were this a major limitation, then in **Fig. 3**, we would have observed many mutants with no defect in gravity but with a defect in LSMMG (e.g., in the top left of the figure). Not only did we not observe a substantial number of such genes, but they were outnumbered by the genes that fell in the bottom right. Therefore, we can conclude that the origination of the library under unit gravity conditions did not impair our ability to investigate this question. Another potential limitation is that analog platforms, such as HARVs, do not completely replicate the space environment and at the gene level there could be numerous differences between these ground-based models and spaceflight not detected within the HARVs^57^. Despite this caveat, numerous studies have demonstrating that physiological changes observed in simulated microgravity can mirror those observed in spaceflight^58–60^, suggesting this approach can be used to explore these global changes in expression.

We provide a comparison of our INSeq data with previously published RNA-Seq data. As shown in **Fig. 6**, there is little correlation between gene requirement (INSeq) and gene expression (RNA-Seq). Although we acknowledge that the INSeq assays do not validate the genome-wide absence of the LSMMG-specific genes or host-associated fitness during animal colonization, it is important to consider both gene requirement and gene expression, as genes that are not induced in a condition may nonetheless be required for growth and/or survival. For example, a gene that is expressed at a constant level under two conditions may be required for survival in only one of those conditions. A broader analysis comparing gene requirement and expression has demonstrated that these categories are often unlinked^61^. Our results argue that genes that are induced in simulated microgravity are not preferentially required for bacterial growth under these conditions.

Overall, this work identifies genes required under exposure to LSMMG, unit gravity, or the general conditions within the analog HARV environment. Importantly, these results demonstrate that simulated microgravity-induced transcripts do not predict which genes are conditionally essential for survival in this environment. Finally, we demonstrated that the HARV environment is amenable to high-throughput genetic experiments and was reproducible within treatments. By laying this groundwork, our findings enable deeper exploration of the genetic mechanisms underlying how microbes adapt and thrive during spaceflight. Such insights are crucial for maintaining healthy host-microbe associations and advancing space biomanufacturing, since enabling reliable biological production processes in reduced gravity is key to optimizing resource utilization and supporting long-term human missions beyond low Earth orbit.

## MATERIALS AND METHODS

### Media and growth conditions

*V. fischeri* strains (**Table 1**) were grown at 25°C in Luria-Bertani salt (LBS) medium (per liter, 10 g Bacto-tryptone, 5 g yeast extract and 20 g NaCl, 50 ml 1 M Tris buffer, pH 7.5, in distilled water). When appropriate, antibiotics or supplements were added to media at the following concentrations: erythromycin, 5 μg/ml; chloramphenicol, 5 μg/ml; X-gal (5-bromo-4-chloro-3-indolyl-β-D-galactopyranoside), 80 μg/ml. Growth media were solidified with 1.5% agar as needed.

**Table 1.**
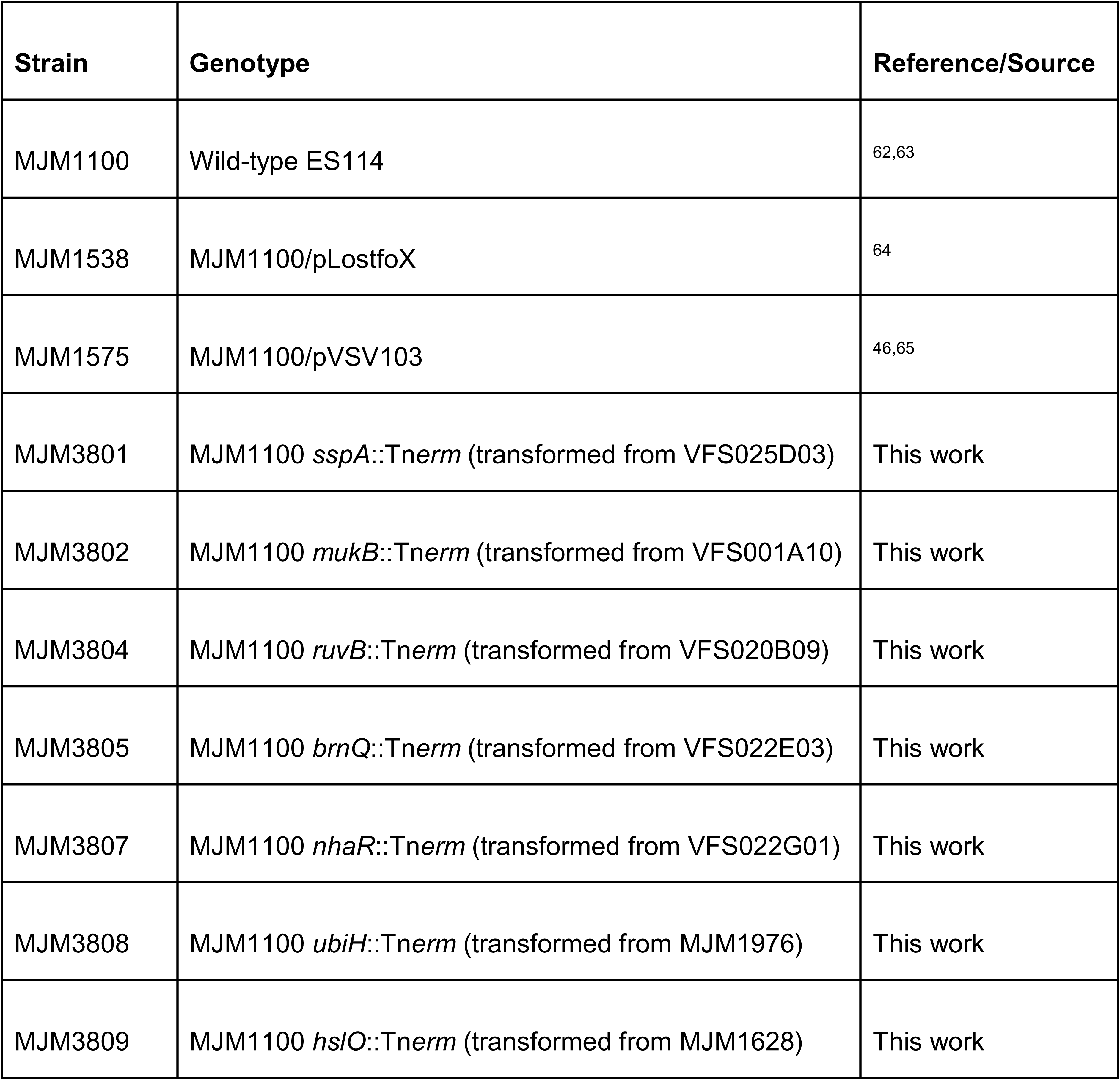
*V. fischeri* strains used in this study.

### Preparation of input and output samples from HARV growth

The Lib04 library of pMarVF1 mariner transposon insertions in MJM1100 (ES114) was described previously^37^. Twenty-five microliters of the library (approximately 2 x 10^8^ CFU) were inoculated into 5 ml LBS, and grown at 26 °C for approximately 5 h, to an OD_600_ of approximately 0.3. Aliquots (552 μl) of culture was transferred to each of eight flasks containing 100 ml LBS (i.e., 2^7.5^-fold dilution) and mixed well; an additional 500 μl of input culture was frozen in 17% glycerol (v/v) as the input sample. From each flask, 50 ml was inoculated into each HARV: LSMMG vessels A, B, C, and D; and gravity vessels A, B, C, and D. Each vessel D was used to monitor growth rate. Samples were grown for approximately 5.25 h to an OD_600_ ≤ 0.3; diluted another 2^7.5^-fold; grown to a final OD_600_ of approximately 0.3, then frozen as the 15-generation output samples.

### INSeq sample preparation

DNA from the input and output samples was prepared using the MoBio Biofilm Isolation Kit (Carlsbad, CA). Samples were prepared for INSeq analysis using the protocol of Goodman *et al.*^45^, with double the BioSamA primer concentration. Samples were submitted to the Tufts University Core Facility (TUCF) for sequencing on the Illumina HiSeq 2500 (single-end 50 bp reads). The resulting reads were deposited at NCBI SRA under accession number SRR12394639.

### Bioinformatic Analysis of INSeq data

Each sample was processed using the bioinformatic software pyinseq (https://github.com/mjmlab/pyinseq, v0.2.0) to quantify transposon insertions and then analyzed using Python visualization and statistical modules. Pyinseq starts by demultiplexing the raw reads using a barcode index and then maps them to the reference genome (ES114v2: CP000020.2, CP000021.2, CP000022.1) using the short-read aligner software Bowtie^66^ with parameters that allow a single base-pair mismatch. Reads with multiple alignments (e.g., to the 12 semi-redundant rRNA operons and in tRNA genes) were excluded. The output alignment files were used to quantify the frequency of reads at each transposon insertion site (TA dinucleotides, for the mariner transposon). Transposon-insertion sites were analyzed if they contain a minimum of three reads and have reads from both the left and right flanking sequence (with a maximum difference of 10-fold abundance for one side over the other). For each sample, a T50 value was calculated, which is defined as the minimum number of transposon insertion sites that account for 50% of the reads in that sample. Gene-level analysis consolidates the site-level data for insertions that fall in the 5’-most 90% of each gene (-d parameter of 0.9). The pyinseq summary gene table (**Table S4**) was further analyzed using pandas, a python module for manipulating large datasets ^67,68^. In addition, pandas was used to calculate spearman correlation values of each pairwise sample comparison. The technical averages for each sample was further analyzed using DESeq2, an R package that normalizes the dataset and performs appropriate statistical tests on high-throughput count data that does not follow a normal distribution such as INSeq^69^. DESeq2 was used to estimate the library size, normalize sequencing depth of samples, and calculate variation of each gene for statistical testing. All *p*-values reported for INSeq data were adjusted for false discovery rate (FDR) using the Benjamini–Hochberg procedure, with genes considered significantly enriched or depleted at an *p*-adj < 0.05.

### Transformation of VFS into MJM1100 background

The VFS mutant collection was assembled by sequencing random pEVS170 transposon insertions into ES114^37,70^. Mutant alleles were moved into the MJM1100 background using transformation under *tfox* induction^71^. Selected VFS strains grown on LBS plates were verified with PCR amplification using locus-specific primers and transposon-anchored primers MJM-127 (pEVS170 transposon) or SamA (pMarVF1 transposon) (**Table S5**). Recipient strain MJM1538 (MJM1100 carrying *pLostfoX)* was grown overnight in 3 ml of LBS containing 2.5 µg/ml chloramphenicol and then subcultured 1:100 into 3 ml of Tris Minimal with *N-*acetylglucosamine (GlcNac) for overnight growth. The recipient was then subcultured 1:50 into fresh Tris-Minimal-GlcNac-Cam with aeration until the OD_600_ reached 0.2-0.3, when 500 µl of recipient culture was incubated with 2.4 µg of VFS donor DNA (prepared with the Qiagen Blood and Tissue kit, Gram-negative bacteria protocol), followed by a brief vortex, and then static incubation at room temperature for 30 min. One ml of LBS was added to culture, transferred to a glass culture tube, and incubated overnight with aeration. The culture was spun down (8000 x *g*, 1 min), 900 μl of supernatant was removed, and the pellet was resuspended in the remaining ∼100 µl of LBS. Aliquots (50 µl) of each sample were plated on LBS-erythromycin (LBS-Erm; 5 μg/ml) and three candidates were selected.

Colonies were restreaked on LBS-Erm plates and then patched on selective media to check for absence of *pLostfoX* (LBS-Cam^S^) and the presence of the transformed DNA (LBS-Erm^R^). Transformation was verified with PCR amplification using primers that target the transposon junction and the gene target (**Table S5**).

### Competitions of individual gene mutants with a marked parental strain in the HARVs

The competitive fitness of seven transposon mutants (**Table 1**) was evaluated in one-on-one competition assays against the LacZ-expressing wild type, MJM1575, in the HARV vessels. Overnight cultures of each strain were prepared in LBS with shaking at 25 °C and diluted 1:100 in fresh medium the following morning. After 2 h of growth at 25 °C with shaking, the mutant and wild-type subcultures were normalized by OD_600_ and combined at a 1:1 ratio in fresh LBS. The resultant starting culture was subsequently loaded into the HARV vessel. At this time, two samples of the mixed input culture were collected: the first was preserved in 33% glycerol (v/v) and stored at -80 °C for later analysis, whereas the second was serially diluted in PBS (pH 7.0) and plated on LBS-Xgal agar (2 mg/ml) to reveal the starting ratio of mutant:control, based on the proportion of white-to-blue colonies, as previously described^37^. In the HARVs, the mutant and marked wild type were grown in competition with each other under LSMMG or gravity conditions for approximately 10 generations at 25 °C and 13 rpm. After 10 doublings, which was determined by OD_600_, samples of the output cultures were preserved in 33% glycerol (v/v) and stored at -80 °C. The input and output samples were plated on LBS-Xgal agar to calculate the competitive index for each sample. The competitive index is equal to the Log_10_ value of the mutant/wild type ratio after competition normalized to its measured ratio at the beginning of the competition.

## DATA AVAILABILITY

Illumina data for the INSeq reads are available at NCBI SRA, Accession number SRR12394639.

## ACKNOWLEDGMENTS

We acknowledge Cheryl Whistler and Randi Foxall for VFS Library mutants and A. Murat Eren and his laboratory for technical advice and mentorship. Funding was provided by NASA’s Illinois Space Grant Consortium Research Seed Grant (M.J.M.), NIGMS R35 GM119627 (M.J.M.), an American Society for Microbiology Undergraduate Research Fellowship (E.B.), and the McNair Scholars Program (E.B.). The project was also supported in part by a NASA Space Biology award 80NSSC18K1465 (J.S.F.). The funders played no role in study design, data collection, analysis and interpretation of data, or the writing of this manuscript.

## AUTHOR CONTRIBUTIONS

EB analyzed the data and wrote the manuscript. MMV conducted the HARV experiments. ER and MMB conducted the INSeq library preparation. MJM and JSF conceived of the study, obtained funding for the study, and wrote and edited the manuscript.

## SUPPLEMENTAL DATA

The file Supplemental_Tables.xlsx includes the following tables:

Table S1: INSeq sample details

Table S2: DESeq2 output for the volcano plots in Figure 4

Table S3: Summary gene table output from pyinseq

Table S4: List of genes depleted as shown in Figure 3

Table S5: Oligonucleotide primers used in this study

