## Supplementary material for "Genome-wide fitness profiling reveals limited modeled microgravity-specific gene requirements in *Vibrio fischeri*": Figure S1

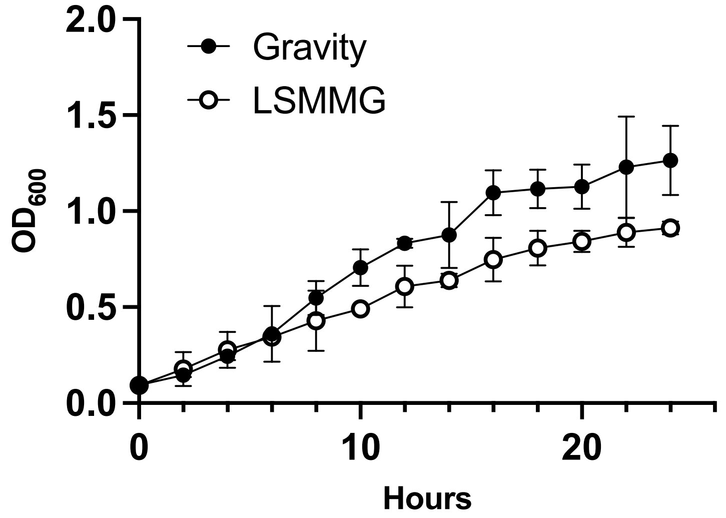


**Figure S1. Growth curves of *Vibrio fischeri* ES114 in the HARV.** Triplicate biological replicates were grown under gravity and low-shear modeled microgravity (LSMMG). Mean and 95% confidence interval are plotted.
